# An Open-source Tool for Analysis and Automatic Identification of Dendritic Spines Using Machine Learning

**DOI:** 10.1101/281667

**Authors:** Michael S Smirnov, Tavita R Garrett, Ryohei Yasuda

## Abstract

Synaptic plasticity, the cellular basis for learning and memory, is mediated by a complex biochemical network of signaling proteins. These proteins are compartmentalized in dendritic spines, the tiny, bulbous, post-synaptic structures found on neuronal dendrites. The ability to screen a high number of molecular targets for their effect on dendritic spine structural plasticity will require a high-throughput imaging system capable of stimulating and monitoring hundreds of dendritic spines in various conditions. For this purpose, we present a program capable of automatically identifying dendritic spines in live, fluorescent tissue. Our software relies on a machine learning approach to minimize any need for parameter tuning from the user. Custom thresholding and binarization functions serve to “clean” fluorescent images, and a neural network is trained using features based on the relative shape of the spine perimeter and its corresponding dendritic backbone. Our algorithm is rapid, flexible, has over 90% accuracy in spine detection, and bundled with our user-friendly, open-source, MATLAB-based software package for spine analysis.

## Introduction

Structural changes in dendritic spines, the tiny postsynaptic protrusions on the dendritic surface of neurons, are considered to be the basis of synaptic plasticity [1] and are known to be important for learning and memory [2]. Dysfunctions in synaptic plasticity are a feature of affective disorders, neurodegenerative diseases, and aging-associated cognitive decline [1].

Recent advances in photostimulation and imaging techniques have made it possible to visualize the morphological and molecular changes in individual spines with high time resolution. Two-photon laser-scanning microscopy in live brain tissue is often used due to its relatively low scattering and precise localization in deep samples [3]. Furthermore, two-photon microscopy can be combined with glutamate uncaging, resulting in targeted photoactivation and plasticity in individual dendritic spines [4]. However, the process of finding, imaging, and analyzing changes in individual dendritic spines is cumbersome and time-consuming. Therefore, the identification of dendritic spines needs to be automated.

Recently, several approaches to semi-automated identification and analysis of dendritic spines have been described [5-10]. These methods have the potential to greatly reduce the amount of effort required for large-scale spine counting and analysis, but are often optimized to a specific cell type, imaging technique, or magnification. Since the majority of spine segmentation algorithms are designed to be used for post-hoc analysis rather than to assist with live imaging, they may require large amounts of computing time and always rely on human input for error correction. Furthermore, variations in image intensity, background signal, and spine length must be accounted for by manual optimization of program settings. Therefore, the application of these algorithms to assist with live spine imaging under varying physiological conditions proves prohibitively difficult.

To reduce errors due to sample variability, some spine identification techniques incorporate machine learning techniques [5, 11]. Since differences in microscopes, fluorescent markers, and spine morphologies lead to variability in how spines are visualized, complex machine learning algorithms such as neural networks and deep learning often require enormous amounts of labeled training images (>10,000), while simpler classifier techniques lack the ability to properly capture the amount of features required to identify spines.

Here we provide a user-friendly tool to analyze, label, segment, and automatically identify dendritic spines. We use a machine learning approach to dendritic spine identification which is highly adaptable to any fluorescent imaging setup. By using adaptive thresholding, we identify neuronal dendrites regardless of background noise and signal intensity. Next, we train a neural network to identify spines based on the position of perimeter pixels relative to the dendrite and spine backbone, as well as the fluorescence intensity along the spine backbone. Our approach is fast and works with a training data set of as few as two thousand images which can be labeled within a few hours using our semi-automated labeling software. Furthermore, our software can be easily adapted to unique imaging setups, and is freely available in open-source MATLAB code.

## Image Acquisition

### Tissue Preparation

To create an algorithm able to detect dendritic spines within a variety of morphologies, the images used for analysis were collected from a variety of genotypes. Organotypic hippocampal slice cultures were prepared as described previously [12] from p4-p6 mice were cultured for 10-12 days before transfection. A biolostic particle delivery system (Helios^®^ Gene Gun System, Bio-Rad) was used to introduce fluorescent GFP labels to obtain sparse transfection of neurons. Two to six days after transfection, neurons in sparsely GFP-labeled CA1 hippocampal regions were chosen for imaging. Individual spines in the striatum radiatum on secondary apical dendrites were chosen for observation.

### Animals

Wild-type C57BL/6J were purchased from Charles river laboratories, and conditional knockout (cKO) lines were generated for IGF1 Receptor and Insulin Receptors as using standard knockout techniques. P4-p6 pups were taken from mothers housed individually in Tecniplast^®^ ventilated cages. Animals were housed on a 12 hour light cycle with a room temperature of 74°F, 50% humidity, with Harlan 7092 ¼” corn cob bedding. All animal procedures were approved by the Max Planck Florida Institute for Neuroscience Animal Care and Use Committee, in accordance with guidelines by the US National Institutes of Health. Max Planck Florida Institute has been AAALAC Accredited since June, 2014.

### Microscopy

Imaging was done on a custom built, two-photon microscope controlled by Scanimage and modified to allow for automated, multiposition image collection [13, 14]. Dendritic spines were imaged over ∼1 hour using a 60X objective and 30X galvanometer-scan zoom (image field ∼8×8µm). One 5µm Z-stack was collected over five Z-planes at each imaging position per minute. Each image was acquired at 128x128 pixels, resulting in a resolution of ∼ 15 pixels per µm in both X and Y.

## Image analysis

The image processing workflow for feature extraction is illustrated in **Error! Reference source not found**.. First, spine locations are labeled in each image, and images are automatically segmented. After segmentation, individual feature vectors consisting of 221 values were used to train a neural network using a scaled conjugate gradient propagation algorithm [15]. Once trained, the neural network was used to evaluate whether feature vectors from newly segmented images represent spine or non-spine locations. All code was written in MATLAB and is freely available at https://github.com/mikeusru/Braintown.

### Preprocessing and binarization

Once an image is loaded (<u>Figure 2A</u>), pixels are converted to grayscale floating-point numbers ranging between 0 to 1. Noise is removed using a standard 2D median filter. Highest-probability background is identified using Otsu’s method of globally thresholding [16]. To ensure no relevant pixels are lost, the global threshold value is reduced by 70%. The average pixel below the background threshold is then subtracted from the image. Next, an adaptive image threshold is computed using local first-order statistics with a neighborhood size of 10×10 µm. Any resulting holes smaller than 0.5µm^2^ are filled. Ideally, the resulting binary image (<u>Figure 2B</u>) includes only regions of neuronal tissue.

**Figure 1:**
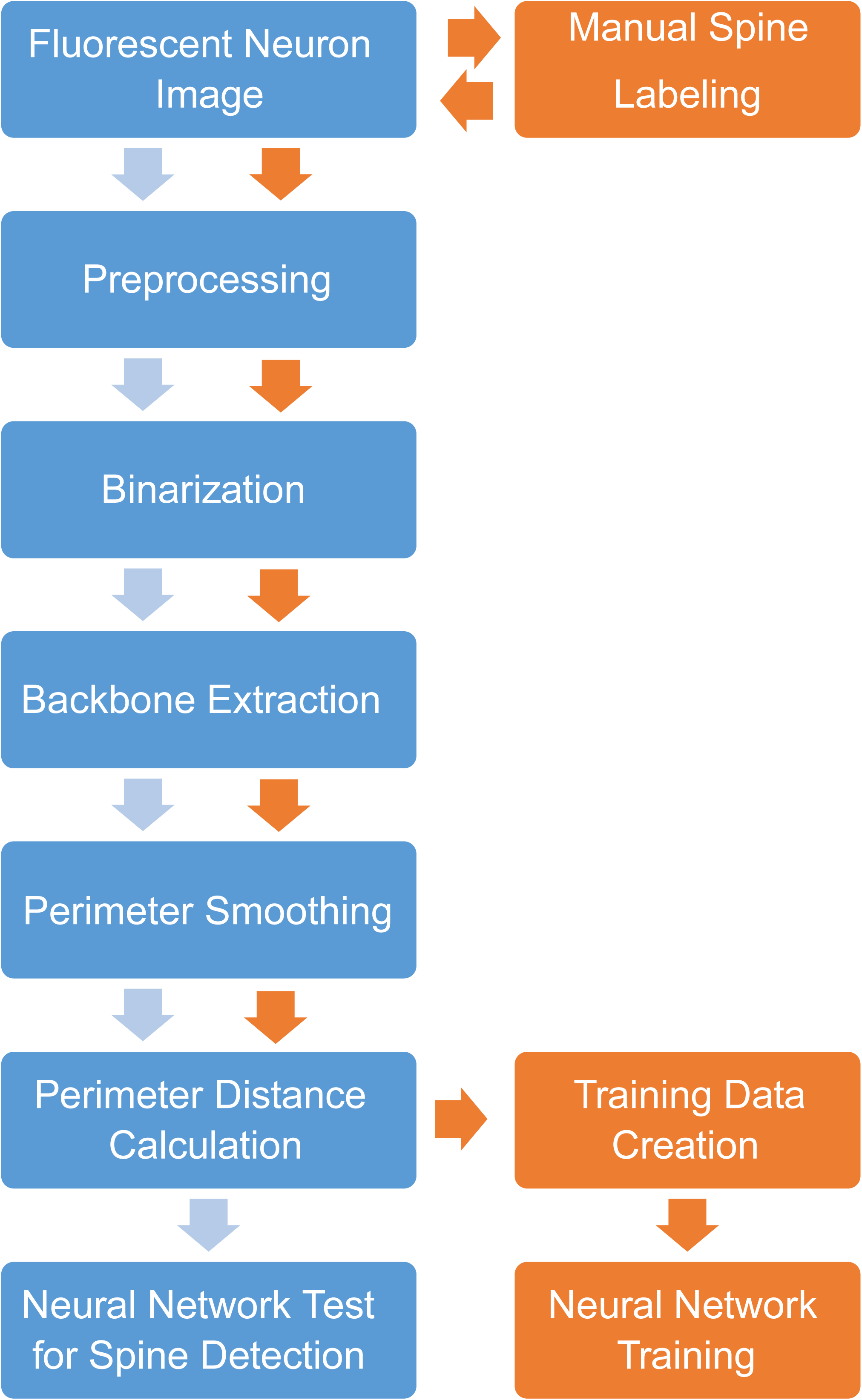
Image processing workflow for automated identification of dendritic spines. Orange: Spine locations are labeled in each image prior to automated segmentation. Extracted feature vectors are used to train a neural network using a scaled conjugate gradient propagation algorithm. Blue: Novel images are preprocessed, segmented, and feature vectors are extracted. Feature vectors are used to evaluate identify potential dendritic spines using the previously trained neural network.

**Figure 2:**
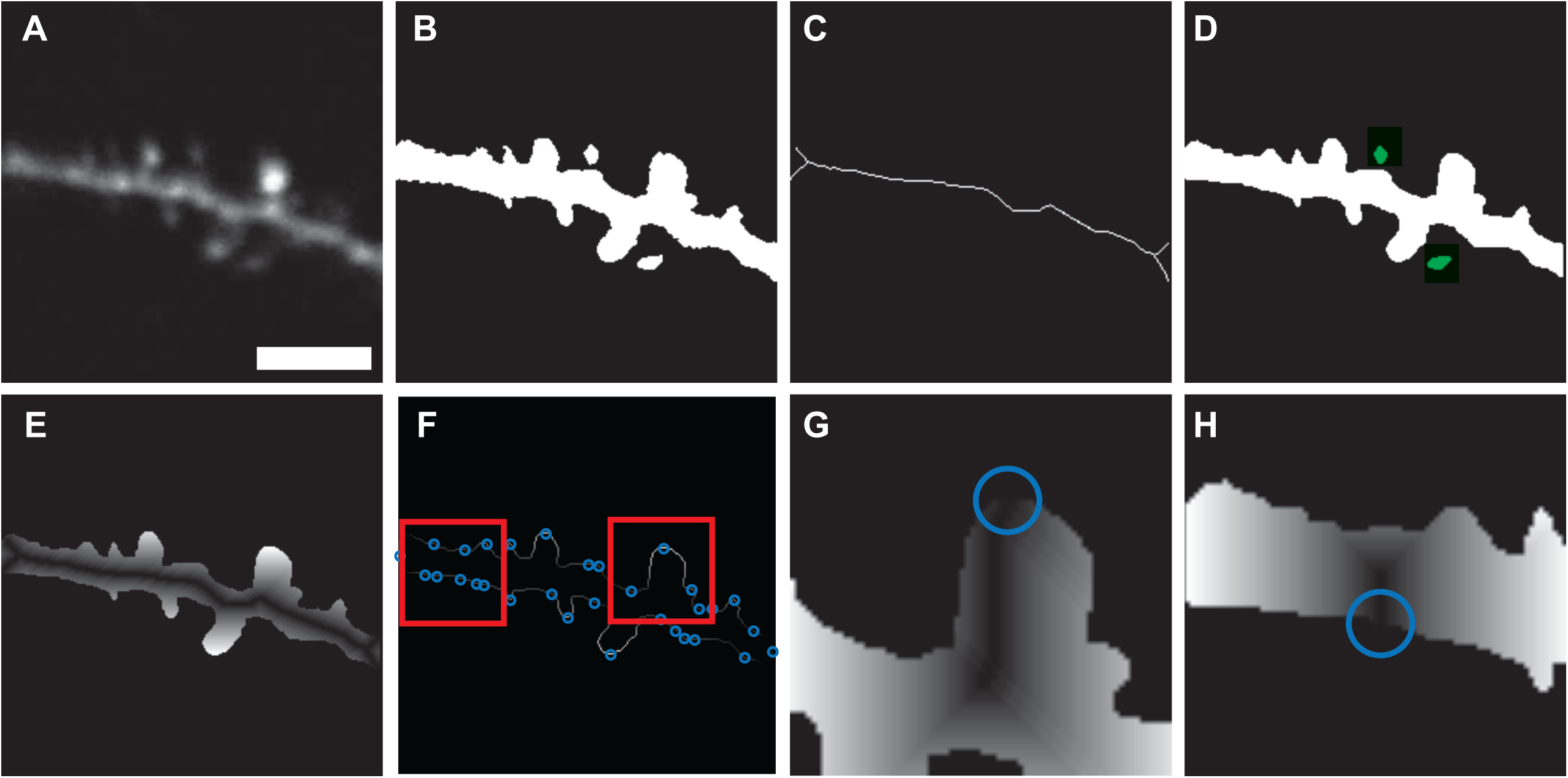
Steps in image segmentation. A. Thresholding using Otsu’s global method, followed by adaptive thresholding. B. Binarization. C. Backbone Extraction D. Identification of removed spines. E. Geodesic distance transform using dendrite backbone as seed location F. Identification of potential spine locations by local maxima along perimeter. G, H. Local geodesic distance transforms for each individual potential spine point, using spine backbone (shortest path between local maxima and dendrite backbone) as seed point. Scale bar = 5µm.

### Backbone extraction

The backbone of the dendrite <u>(Figure 2C</u>) was identified by thinning the binary image until all structures had a thickness of no more than one pixel, and then removing any branches which didn’t belong to the dendrite. After skeletonization [17], the presence of dendritic spines and noise within the image causes a significant number of spurious branches and loops which are not representative of the dendrite itself. Loop artifacts were removed by filling in all areas smaller than 0.5µm^2^ and undergoing a second round of skeletonization. Any isolated segments where p<M were removed, where p was the amount of pixels in the segment, and M was the maximum spine length (2µm) multiplied by the number of pixels per µm. The remainder of the spurious segments were removed by a recursive trimming algorithm adapted from Cheng et al. [7]. Basically, endpoint pixels were iteratively removed from the skeleton and added to a set of deleting templates through the use of a nested loop. If the iteration did not add to the deleting template, then the deleting template was permanently removed from the skeleton. The code structure is presented below:

1. **Initialize m = 1**
2. **Repeat until m = M**
  a. **Initialize removed segments = blank**
  b. **Repeat m times:**
    i. **Find skeleton endpoints, ignoring those near border**
    ii. **Add skeleton endpoints to removed segments**
  c. **Remove skeleton endpoints from skeleton**
  d. **Restore any removed segments that have m pixels**
  e. **m = m + 1**

After trimming, the backbone often retained some small kinks leftover from the initial skeletonization process. As these kinks could introduce artifacts in the later perimeter distance calculation, they were removed by a custom smoothing algorithm also adapted from Cheng at al. [7]: First, all branch points belonging to the initial, untrimmed skeleton were located along the dendrite backbone. Next, the branchpoints were dilated by M/4 to include all local backbone pieces which might belong to a kink. Finally, these kinks were removed, and the resulting line endpoints connected, resulting in a smooth backbone segment.

### Surface smoothing

To isolate individual segments of the cell perimeter to be used as features for spine detection, the surface of the binary object needed to be smooth, lacking any spurious pixels or diagonally connected regions. Smoothing was achieved using an array of morphological operations on the binary image. First, a majority operation [18] set a pixel to 1 if five or more pixels in its 3x3 neighborhood are 1s, otherwise the pixel is set to 0. Next, the image is morphologically opened, closed, and opened again using a 3x3 structuring element of ones. Pixels connected to fewer than three other pixels were removed, and a diagonal fill was used to eliminate any 8-connectivity of the background, essentially transforming diagonal connections into right angles. The binary objects were then thickened by adding a one-pixel width border, as long as that border did not form a new connection with a neighboring border. An example of the result attained through surface smoothing can be seen in <u>Figure 3</u>.

**Figure 3:**
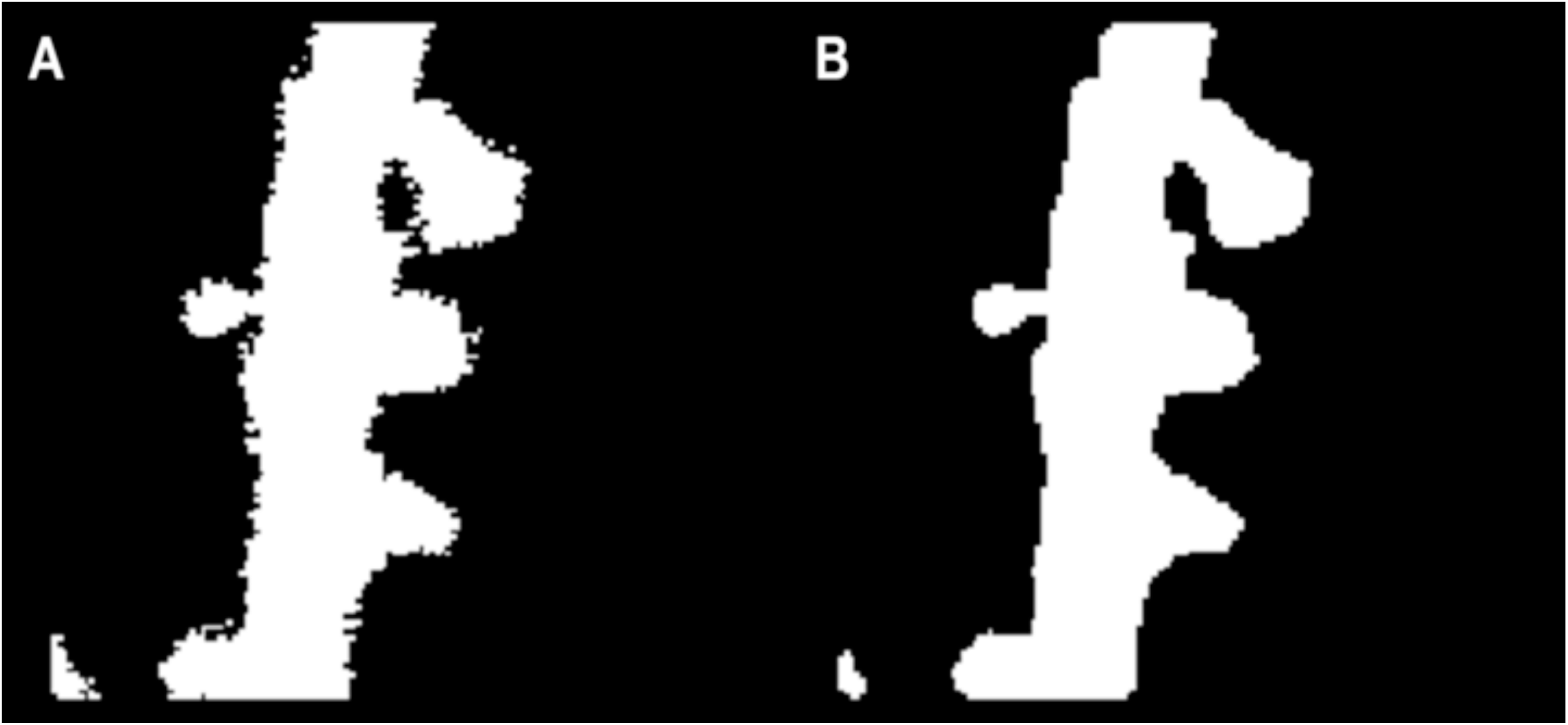
Surface smoothing of binary image. (A) Initial binary image contains a significant noise around its perimeter. (B) Surface-smoothed image lacks kinks, as well as spurious or diagonally – connected pixels.

### Identification of disconnected spines

As the purpose of our algorithm was to find spines which were obviously attached to the dendrite, small objects which became disconnected from the dendrite <u>(Figure 2D</u>) during the surface smoothing step were categorized using k-means clustering, but were ignored from the neural network training data. In any individual image, the signal to noise ratio was calculated in objects that were within distance M from the dendrite backbone. Each respective signal was collected from the pixels in the original image <u>(Figure 2A</u>) which overlapped with the object, while noise was calculated using pixels in the bounding box of the object minus the pixels within the object. If more than two objects were detected, spines were identified using k-means clustering of the signal to noise ratios. All objects detached from the main dendrite structure were ignored for the remaining calculations.

### Perimeter feature extraction

Three feature vectors were used for neural network training and spine identification: perimeter distance from dendrite backbone (PD), perimeter distance from spine backbone (PS), and fluorescence intensity along the spine backbone (IS). The location of each feature vector, as well as the individual values of PD features, were quantified based on a geodesic distance transform [19] of the binary image of the dendrite, using the dendrite backbone as a seed location. Thus, the value assigned to each connected pixel represents its relative distance from the dendrite backbone (<u>Figure 2E</u>). The central position of each feature vector was assigned by finding local maxima along the perimeter of the geodesic transform (<u>Figure 2F</u>), and a geodesic distance transform of the perimeter itself (<u>Figure 4A</u>) served to organize all perimeter pixels into relative locations. Thus, each PD feature vector represents a 5µm segment of pixel values along the edge of the geodesic transform (<u>Figure 4C</u>).

**Figure 4:**
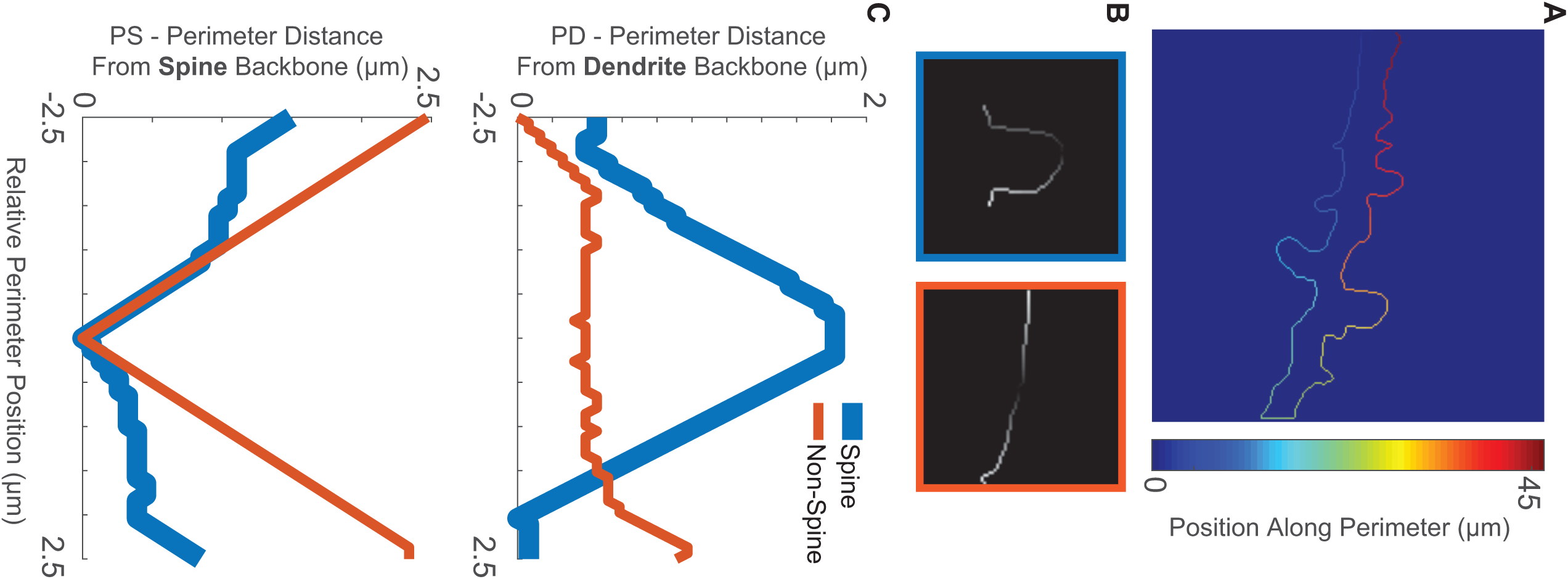
PD and PS feature vectors. (A) Relative position values, in µm, assigned to each pixel starting from a random seed and following a single direction. (B) 5µm perimeter segments are extracted at each potential spine point to create PS feature vectors. Brighter pixels indicate higher distance from spine backbone. Left box (blue) is a true spine, right box (orange) is a non-spine. (C) Perimeter feature vectors represent respective pixel values (y axis) and pixel position (x axis).

Unlike PD features, which represent distance from the dendrite backbone, PS features represent distance from the spine backbone. The spine backbone was identified as the shortest path between the feature origin point on the perimeter (<u>Figure 2,G,HF</u>) and its nearest point on the dendrite backbone with the help of a fast marching algorithm [20]. A geodesic distance transforms was calculated using the spine backbone as a seed (<u>Figure 2G,H</u>), and PS features are represented as a 5µm segment of pixel values along the resulting perimeter (<u>Figure 4B,C</u>).

PD and PS features were arranged based on their respective position along the perimeter. To minimize the amount of necessary training data, position information was defined by a single value as the directional distance along the perimeter from the center of the feature origin. To assign position values, a closed-loop perimeter was first cut at a random point. A geodesic distance transform, with one endpoint as a seed, was then used to assign a single value to each pixel <u>(Figure 4A</u>). As a result of the transform, each consecutive pixel was assigned a value based on its travel distance from the seed pixel. For this transform to work properly, it was pertinent that the perimeter lack any kinks or loops, as this would result duplicate position values. By assigning these position values to each perimeter feature, we translated 2D perimeter images (<u>Figure 4B</u>) into 1D arrays of feature-specific values (<u>Figure 4C</u>). Finally, since the amount of pixels in a 5µm segment varied based on the resolution of the initial image, PS and PD feature vectors were standardized by interpolating to 100 values each.

Features in the IS group were assigned using pixel positions from the spine backbone, and pixel values from the original image. The resulting feature vector represents a line of intensity values starting at the dendrite backbone and finishing at the tip of the spine. Due to the spine backbone varying in length, each group of values was interpolated to 20 features <u>(Figure 5A</u>), while the 21^th^ feature represented the original spine backbone length in µm <u>(Figure 5B</u>).

**Figure 5:**
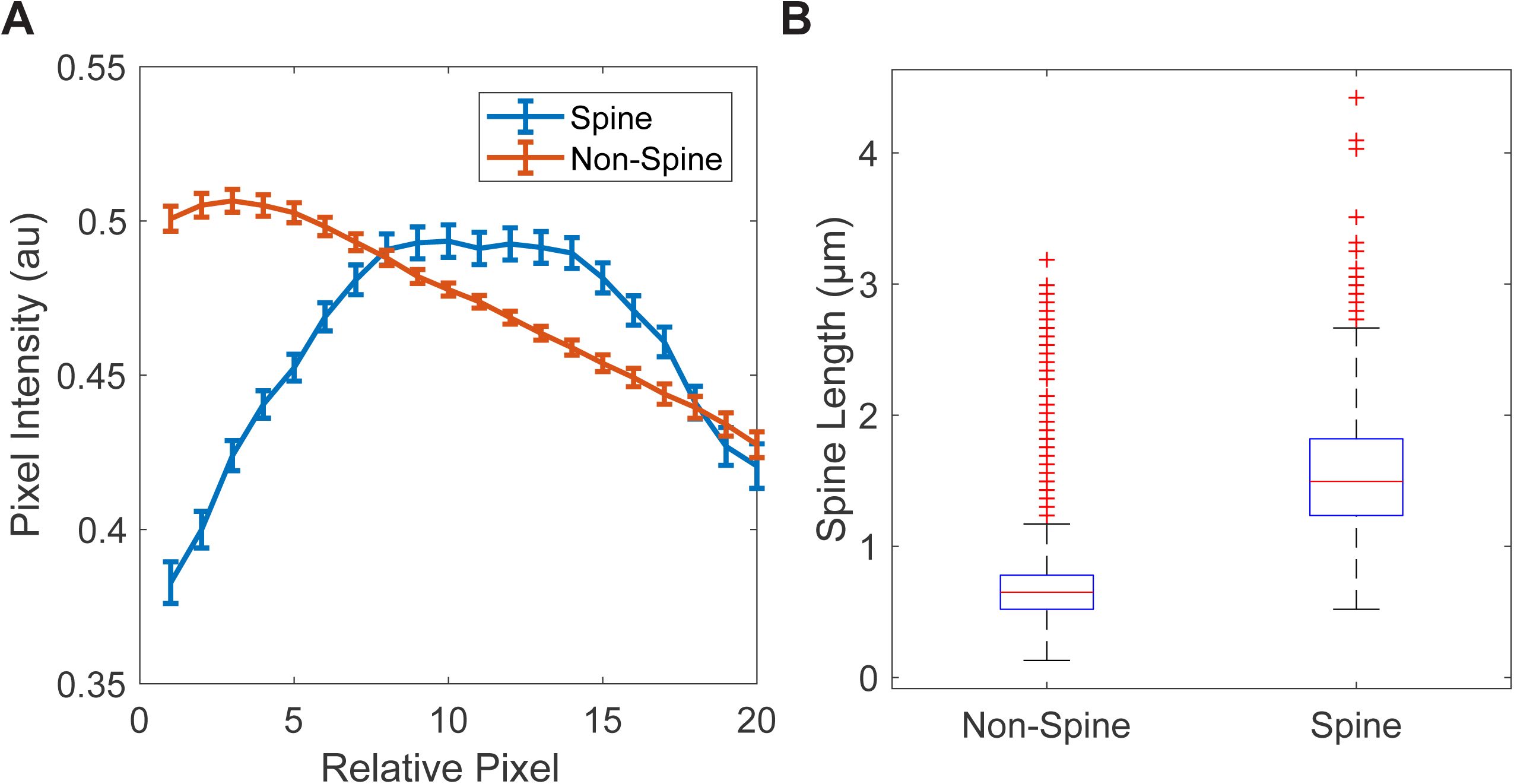
Length and Intensity of the spine backbone. (A) Average pixel intensity in pixels taken along line connecting tip of potential spine to closest point in dendrite backbone. (B) Average length of spine backbone in pre-labeled Spine and Non-Spine objects.

## Network Training

PD, PS, and IS feature sets consisting of a combined 221 values were used to train a neural network using a scaled conjugate gradient propagation algorithm. The network had one hidden layer with 20 nodes, as these parameters showed to elicit the highest accuracy in spine categorization while keeping training and classification times manageable. Features sets were classified as either spine or non-spine. To label training data, we designed an application which allows users to rapidly identify dendritic spines by clicking on their location in an image. A 1×1µm box was then drawn around each identified spine. Boxes that were within 1.5µm from the image border were ignored to avoid edge artifacts. Boxes which overlapped with a disconnected blob (<u>Figure 2</u> **Error Reference source not found**.D) were ignored as well. Feature sets were classified as spines if their point of origin was inside the box.

## Software Design

To make our tools accessible to users who may lack any significant coding expertise, we built a straightforward front-end user interface for viewing, analyzing, labeling, and segmenting images of dendritic spines in MATLAB <u>(Figure 6</u>). The main window (**Error! Reference source not found**.A) allows users to load either individual images or image sets, browse through the loaded data (**Error! Reference source not found**.B), and restrict certain files from being loaded (**Error! Reference source not found**.C). Drop-down menus also let users perform common calculations such as 3D projection on Z stacks and drift correction on timelapse image sets. Users can draw circular or polygonal ROIs on the image (**Error! Reference source not found**.E) to calculate changes in spine volume over time (**Error! Reference source not found**.D). While remaining compatible with variable data sources, this program is particularly tuned to analyze data collected using our automated multiposition imaging system [14]. A semi-automated spine selection tool for labeling training data is also provided (**Error! Reference source not found**.F). Users can enter spine selection mode (**Error! Reference source not found**.G), where clicking on the image frame will label and store the local coordinates of each spine (**Error! Reference source not found**.E). Users have the option to track spines through brightness, where given a timelapse image set, spine coordinates will automatically update to their new closest position. Finally, the spine selection tool allows users to train, preview, and test a neural network for its capability to find dendritic spines (**Error! Reference source not found**.H).

**Figure 6:**
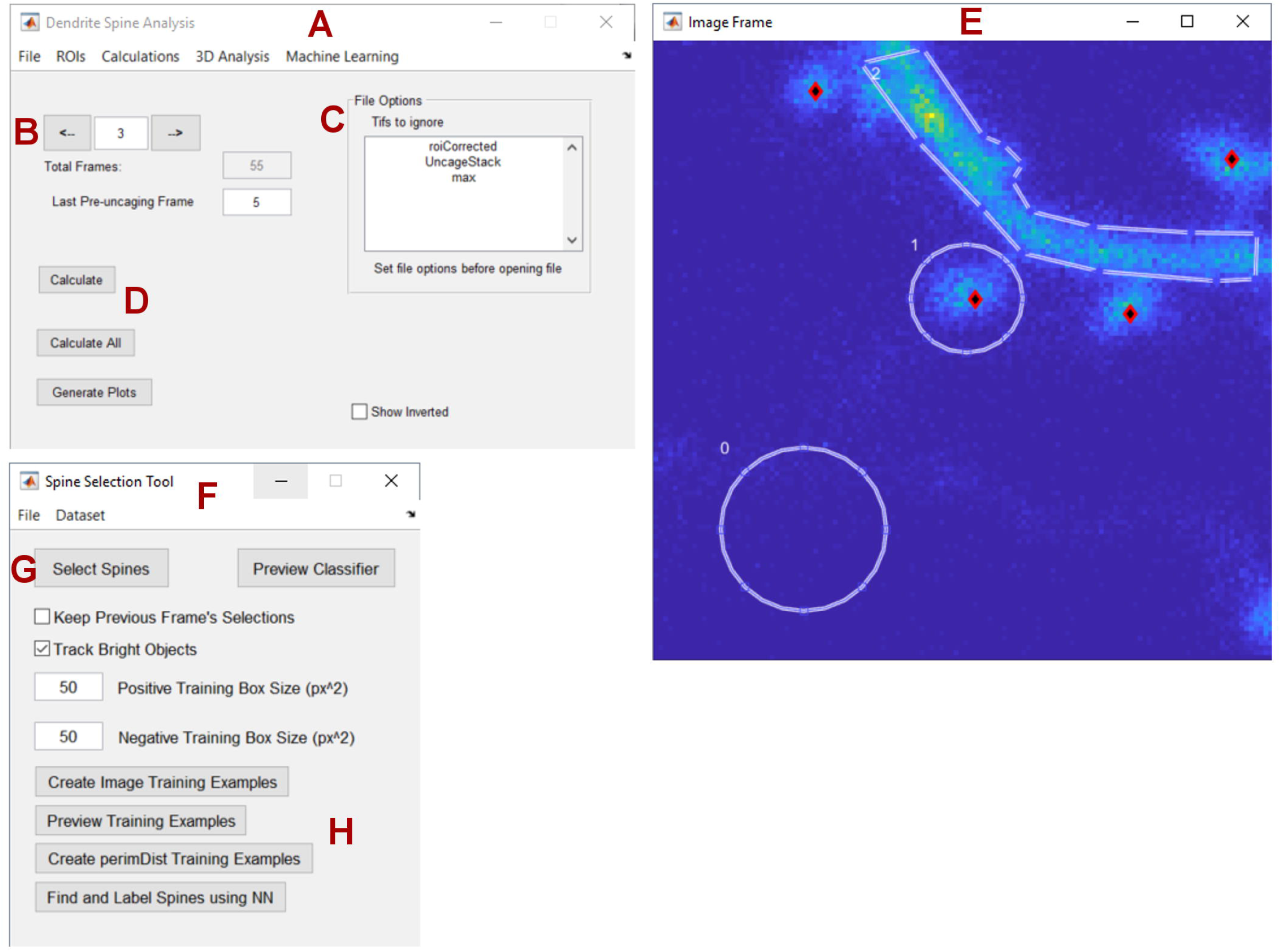
Graphical user interface to label and measure image regions. (A) Base dendrite analysis window responsible of loading/saving data, (B) switching between images, (C) avoid particular files, and (D) running calculations on ROI area over time. (D) Image preview window for drawing polygonal/circle ROIs, and identifying individual spines (red/black rhombi). (F) Spine selection and machine learning tool allows toggling of (G) spine selection mode, semi-automated spine tracking, (H) gathering and previewing of training data, and spine finding using a trained neural network.

To create a powerful yet user-friendly system for image segmentation, we created a modular interface where users can manually select, customize, evaluate, and share plugins and configurations without any coding experience (<u>Figure 7</u>). A function selection window (<u>Figure 7A</u>) loads all of the plugins from a local plugins folder and displays them in an alphabetized list (<u>Figure 7B</u>). Each plugin serves as a step in the image segmentation, analysis, or feature extraction process, and may have unique inputs and outputs. Using drop-down lists, users may select which output variables will serve as inputs for plugins down the line. For example, the function selected in <u>Figure 7C</u> takes the input variable “BW (1)”, and outputs “thin (2)” and “spineSearchZone (2)”, both of which are used as inputs in other steps down the line. A number referencing the analysis step is attached to each variable name to avoid errors where multiple plugins have outputs with the same name. To clarify the types of input and output variables associated with each plugin, as well as the general function of the plugin itself, an informational window previews all relevant information as each function is selected (<u>Figure 7D</u>). Once the custom segmentation process is run, all individual output variables are previewed as images in a separate window (<u>Figure 7E</u>). Once users are satisfied with their plugin configuration, the configuration can be saved, shared, commented on, and even rated for success at a certain task by multiple users (<u>Figure 7F</u>).

**Figure 7:**
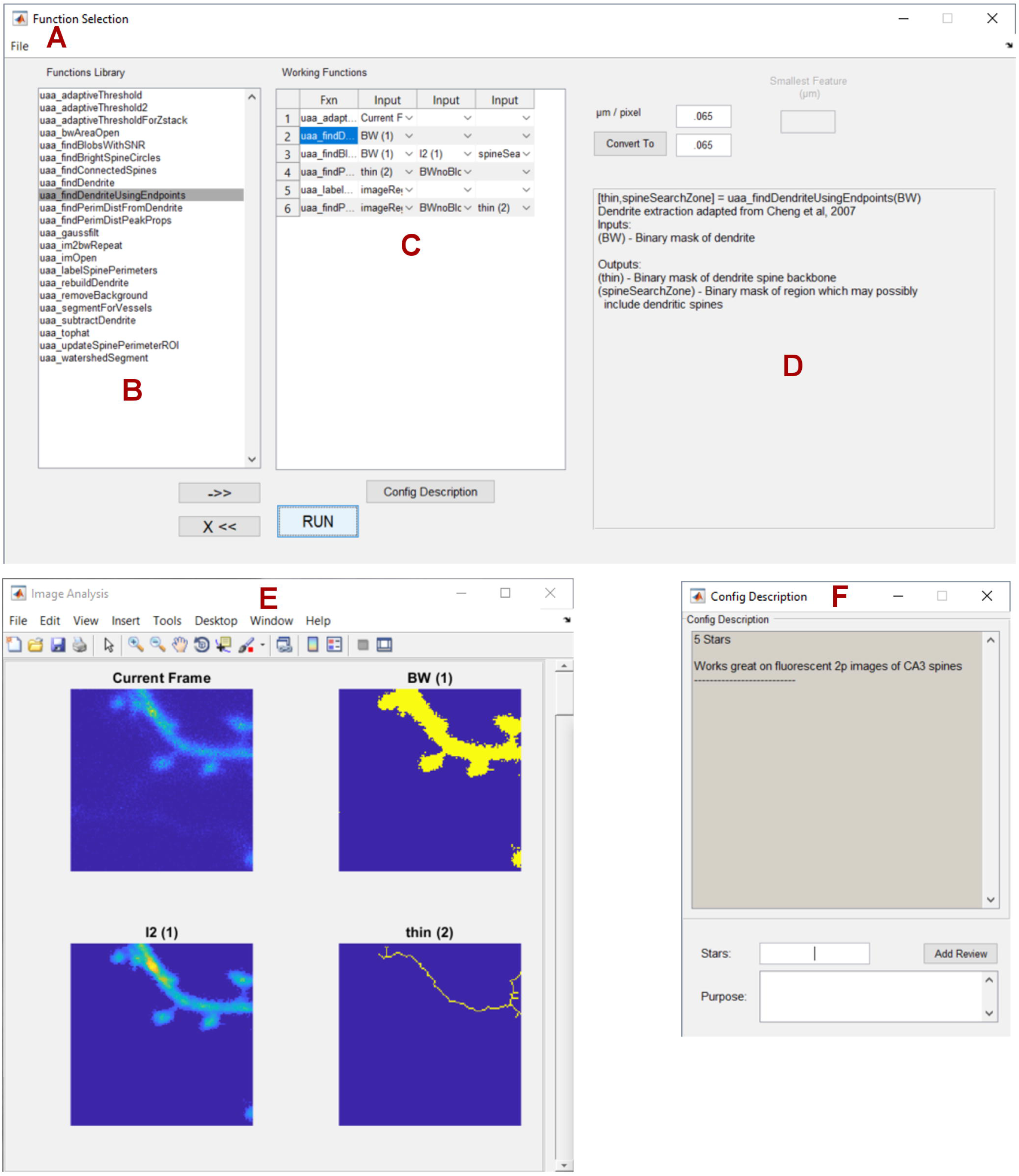
Graphical user interface for image segmentation. (A) The function selection window handles the available functions and their configuration. (B) All available functions are displayed in the plugin repository. (C) Selected functions and their inputs are selected in the current configuration space. (D) A short tutorial for each individual plugin is displayed upon selection. (E) Outputs of each plugin are previewed in a separate, scrollable window. (F) Plugin configurations can be saved, shared, and rated between multiple users.

## Results and Discussion

We used 1837 images to train, validate, and test the neural network. 3627 and 11922 feature sets were categorized as spine and non-spine, respectively. Spine PD feature arrays were often marked with a pseudo-linear increase, and then a decrease, indicating the protruding shape of the spine, while non-spine PD arrays tended to be flat, with a lower amplitude at the center (<u>Figure 4C</u>, <u>Figure 8A</u>). PS feature arrays, on the other hand, tended to have lower amplitudes when associated with a spine, and had a more pronounced V-shape at non-spine positions (<u>Figure 4C</u>, <u>Figure 8B</u>).

**Figure 8:**
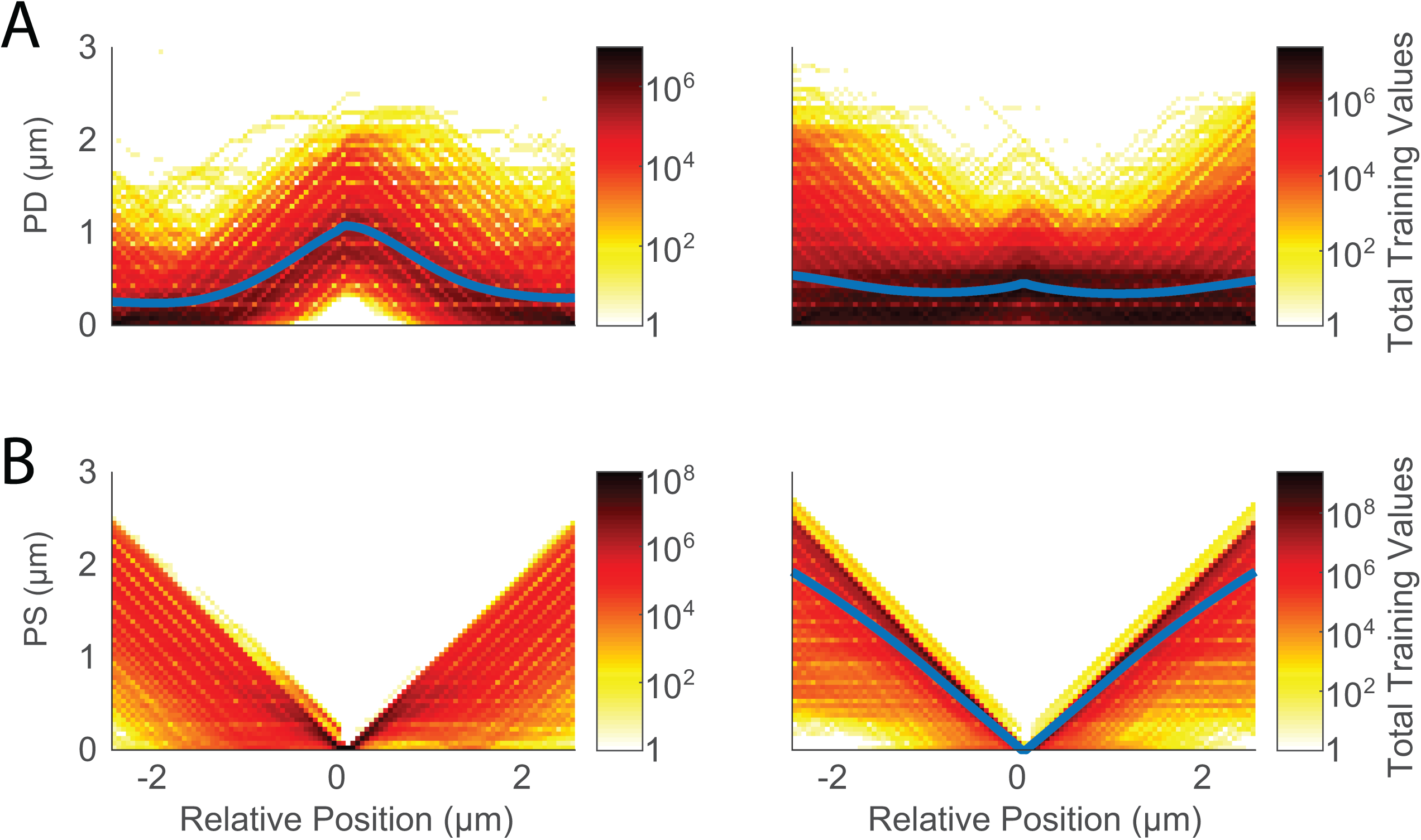
Cumulative PD and PS features. Each chart represents total points binned from all feature vectors in. (A) Left – Spine, Right – Non-Spine binned feature vectors. (B) Left – Spine, Right – Non-Spine binned feature vectors. Blue lines represent the average of all feature vectors in each group.

As expected, the length of the spine backbone tended to be longer in spines versus non-spines <u>(Figure 5B</u>). Spine IS feature arrays often had a pronounced increase followed by decrease in amplitude, indicating the bright center of the dendritic spine, while non-spine groups were characterized by a more linear drop-off in signal <u>(Figure 5A</u>).

Labeled data was split into three groups – Training (60%), Validation (20%), and Testing (20%). Classifier results for training and testing data are shown in <u>Figure 9</u>. Overall, classification accuracy was highly similar between training and test datasets, indicating that there was no overfitting of the model. In the testing dataset, 94.5% of actual spines were classified as spines (true positive), and 5.5% were classified as non-spines (false negative). 98.5% of non-spines were classified as non-spines (true negative) and 1.5% classified as spines (false positive). These results indicate that our model is highly successful in spine identification. The fact that none of the data from the testing group was used to train the model indicates the high predictive value of our algorithms. Furthermore, spine identification in dendrites collected at magnification values different from those of the training data (<u>Figure 9B</u>) predicts the scalability of our algorithm to broader datasets.

**Figure 9:**
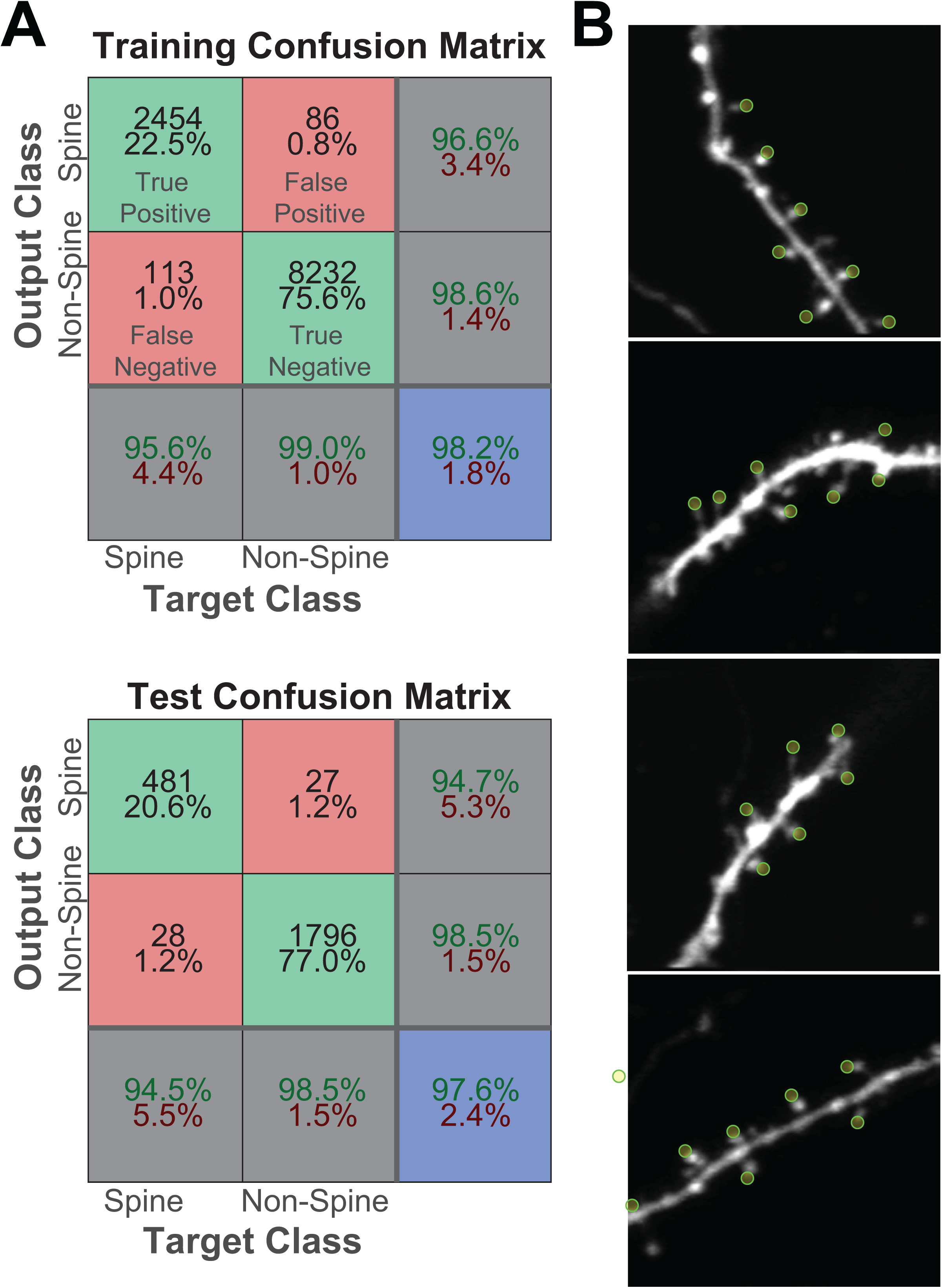
Neural Network’s ability to identify spines. (A) Training and Test confusion matrices. (B) Yellow circles indicate spines identified by neural network in naïve images.

A major benefit of our spine classification algorithm is a lack of parameters which users are required to tune. The only input required by the algorithm is the relative scale of the image in pixels/µm, which can often be extracted automatically from images saved with modern imaging software. Furthermore, we expect our algorithm to become more powerful and accurate for dendritic spine identification as more training data becomes available. Therefore, we see our algorithm as a particularly user-friendly option for those looking to automate fluorescent imaging and/or targeting of dendritic spines. In particular, we expect that the combination of this technique with our previously developed spine imaging automation software [14] will lead to significant increases in the throughput of spine imaging and stimulation.

While many techniques have been developed to identify dendritic spines [5-11, 21], many of these techniques were specifically designed for post-hoc analysis, relying on additional human input to correct mistakes. While our algorithm does not claim to have 100% accuracy, its goal is to identify a large majority of clearly demarcated spines within a sample for automated imaging and photostimulation. For such an automated system to work, spine labeling must require no human input, and have a minimal number of false positives, which lead to throwaway data. Our algorithm accomplishes precisely this feat, relying only on machine learning and previously labeled training data. Furthermore, to minimize the amount of human training time necessary to train the neural network algorithm, we’ve taken steps to simplify our data as much as possible, reducing images to a set of feature vectors which convey important information about spine shape. Since our training data and code is open-sourced and shared online, we expect other labs to build upon and improve our algorithm by adding their own training data, therefore increasing the potential accuracy of spine identification. Furthermore, our software can add additional training features, allowing for even further improvements of detection accuracy.

## Conclusion

Overall, we believe that our neural network model for automated spine identification in fluorescent neurons is highly accurate, scalable, and is built to easily be upgraded with the addition of training data and programmatic improvements. Due to its open-source availability, simplicity, and lack of tunable features, we expect this software to be used both in post-hoc spine analysis, as well as for automated spine tracking during imaging experiments.

